# Antibody responses to SARS-CoV-2 mRNA vaccines are detectable in saliva

**DOI:** 10.1101/2021.03.11.434841

**Authors:** Thomas J. Ketas, Devidas Chaturbhuj, Victor M Cruz-Portillo, Erik Francomano, Encouse Golden, Sharanya Chandrasekhar, Gargi Debnath, Randy Diaz-Tapia, Anila Yasmeen, Wilhem Leconet, Zhen Zhao, Philip J.M. Brouwer, Melissa M. Cushing, Rogier W. Sanders, Albert Cupo, P. J. Klasse, Silvia C. Formenti, John P. Moore

**Affiliations:** Department of Microbiology and Immunology, Weill Cornell Medicine, New York, NY 10065; Department of Radiation Oncology, Weill Cornell Medicine, New York, NY 10065; Department of Urology, Weill Cornell Medicine, New York, NY 10065; Department of Pathology and Laboratory Medicine, Weill Cornell Medicine, New York, NY 10065; Department of Medical Microbiology and Infection Prevention, Amsterdam University Medical Centers, Location AMC, University of Amsterdam, Amsterdam Infection & Immunity Institute, 1105 AZ Amsterdam, the Netherlands; Department of Medicine, Weill Cornell Medicine, New York, NY 10065

**Keywords:** SARS-CoV-2, S-protein, RBD, COVID-19, saliva

## Abstract

Vaccines are critical for curtailing the COVID-19 pandemic (1, 2). In the USA, two highly protective mRNA vaccines are available: BNT162b2 from Pfizer/BioNTech and mRNA-1273 from Moderna (3, 4). These vaccines induce antibodies to the SARS-CoV-2 S-protein, including neutralizing antibodies (NAbs) predominantly directed against the Receptor Binding Domain (RBD) (1-4). Serum NAbs are induced at modest levels within ∼1 week of the first dose, but their titers are strongly boosted by a second dose at 3 (BNT162b2) or 4 weeks (mRNA-1273) (3, 4). SARS-CoV-2 is most commonly transmitted nasally or orally and infects cells in the mucosae of the respiratory and to some extent also the gastrointestinal tract (5). Although serum NAbs may be a correlate of protection against COVID-19, mucosal antibodies might directly prevent or limit virus acquisition by the nasal, oral and conjunctival routes (5). Whether the mRNA vaccines induce mucosal immunity has not been studied. Here, we report that antibodies to the S-protein and its RBD are present in saliva samples from mRNA-vaccinated healthcare workers (HCW). Within 1-2 weeks after their second dose, 37/37 and 8/8 recipients of the Pfizer and Moderna vaccines, respectively, had S-protein IgG antibodies in their saliva, while IgA was detected in a substantial proportion. These observations may be relevant to vaccine-mediated protection from SARS-CoV-2 infection and disease.

---

During December 2020, the availability of the Pfizer and Moderna vaccines provided an opportunity for us to assess the development of antibody responses to the SARS-CoV-2 S-protein and its RBD in serum and saliva samples from immunized HCWs participating in the NYP-WELCOME trial. (Pfizer, Group 1, n =40; Moderna, Group 2, n = 9). For comparison, we used two sub-groups of non-vaccinated individuals who were SARS-CoV-2 uninfected (Group 3, n = 8) or who had recovered from infection during or prior to participation in the trial (Group 4, n = 6). Among the Group-4 members, 3 were vaccinated. Further analyses of Group-4 members, and others who may have become infected during the course of the study, are in progress.

Longitudinal profiles of serum and saliva S-protein IgA, IgG and IgM responses in selected individuals from Groups 1-4 are shown in Fig. 1. Additional longitudinal profiles are shown as Extended Data (ED Fig. 1). The timing of the approximately monthly NYP-WELCOME study visits was not coordinated with the dates on which participants were vaccinated. Hence, samples were not available for some participants in the 3-(Pfizer) and 4-week (Moderna) period between the two vaccine doses. A collated data set for all the vaccinated participants is presented in Fig. 2, which also includes antibody reactivity with the SARS-CoV-2 RBD. The proportions of vaccinated individuals with saliva and serum IgG, IgM and IgA S-protein antibodies after the first and second doses are summarized in Table 1. Both serum IgA and secretory IgA (SIgA) are detected in the ELISA (Methods and ED Fig. 2).

**Table 1.** Proportions of vaccinated individuals with saliva and serum IgG, IgM and IgA S-protein antibodies after the first and second doses. Among the 40 Pfizer and 9 Moderna vaccine recipients, only 13 and 6 provided saliva samples in the period after dose 1 but before dose 2. However, 37 and 8 of them provided saliva samples after dose 2. Two serum samples from Moderna vaccine recipients were not available for analysis.

| Group 1 – Pfizer vaccine |  |  |  |  |  |  |
| --- | --- | --- | --- | --- | --- | --- |
|  | IgA |  | IgG |  | IgM |  |
|  | Saliva | Serum | Saliva | Serum | Saliva | Serum |
| After Dose 1 | 2/13 (15%) | 6/10 (60%) | 3/13 (23%) | 7/10 (70%) | 0/13 (0%) | 2/10 (20%) |
| After Dose 2 | 22/37 (59%) | 36/36 (100%) | 37/37 (100%) | 36/36 (100%) | 8/37 (22%) | 27/36 (75%) |

| Group 2 – Moderna vaccine |  |  |  |  |  |  |
| --- | --- | --- | --- | --- | --- | --- |
|  | IgA |  | IgG |  | IgM |  |
|  | Saliva | Serum | Saliva | Serum | Saliva | Serum |
| After Dose 1 | 5/6 (83%) | 5/5 (100%) | 5/6 (83%) | 5/5 (100%) | 1/6 (17%) | 3/5 (60%) |
| After Dose 2 | 7/8 (88%) | 7/7 (100%) | 8/8 (100%) | 7/7 (100%) | 1/8 (13%) | 4/7 (57%) |

**Figure 1.**
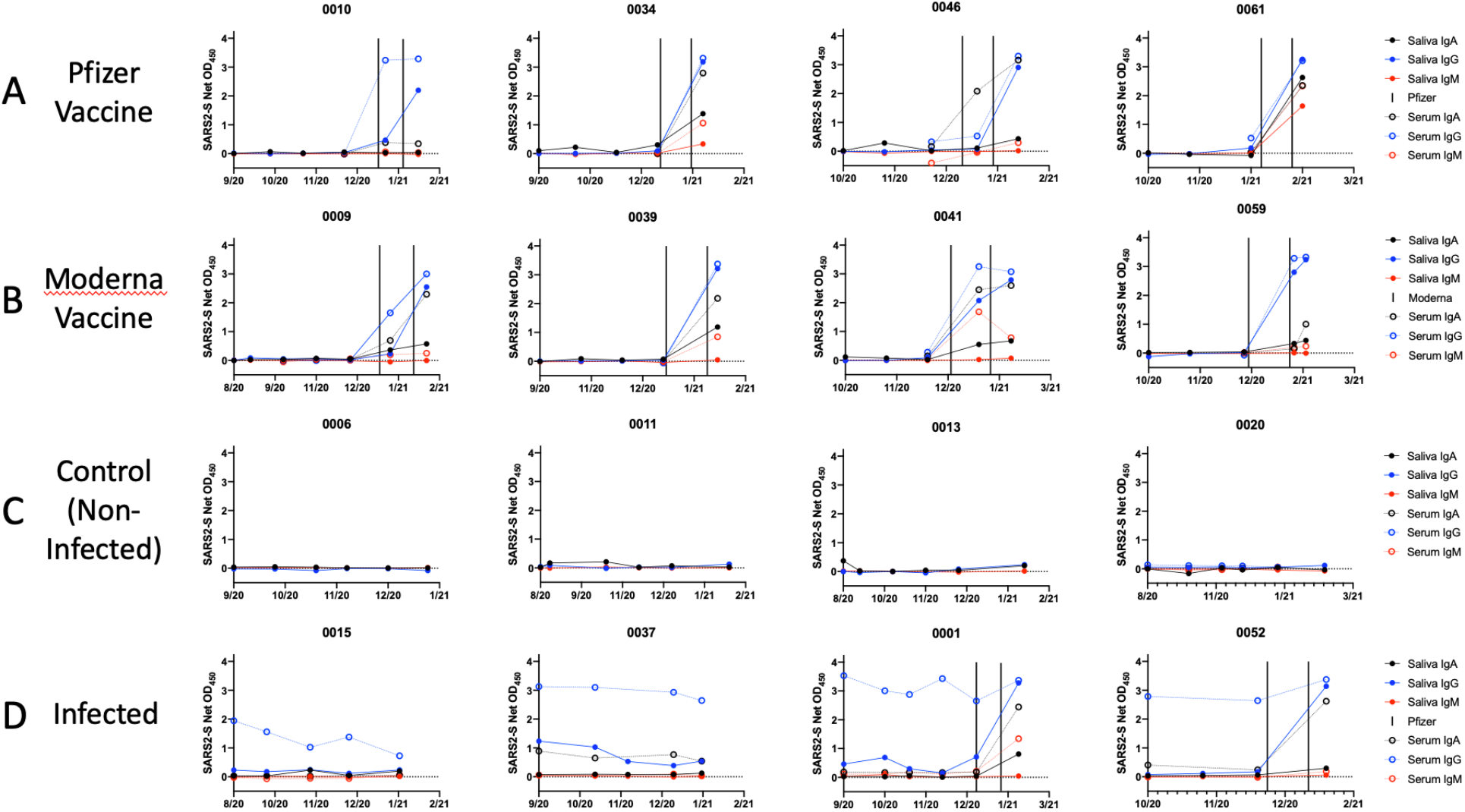
Antibody response to the SARS-CoV-2 S-protein in saliva and sera from SARS-CoV-2 vaccine recipients and infected people. Each diagram shows S-protein IgA, IgG, and IgM antibody reactivities over the time of sampling. The dates of vaccination are indicated by the variable bars. Representative single-dilution binding data are shown for sera from each category: (A) Pfizer vaccine: Cases 0010, 0046, 0061, 0034. (B) Moderna vaccine: Cases 0016, 0078, 0041, 0059. (C) Control (non-infected): Cases 0016, 0011, 0013. 0020. (D) Infected: Cases 0037, 0063, 0001, 0052 (the latter two were vaccinated). Additional profiles are shown in ED Fig. 1.

**Figure 2.**
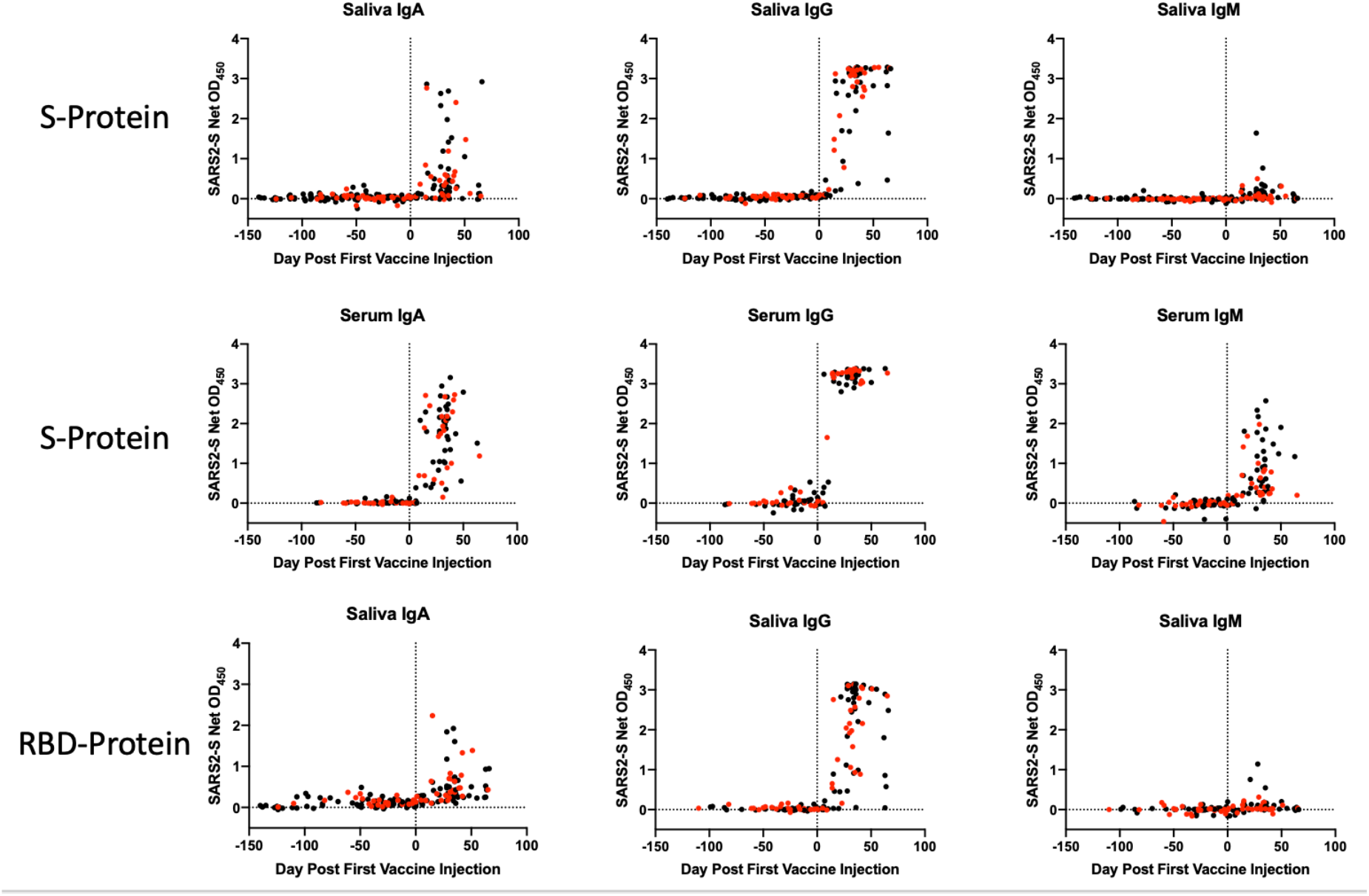
Antibody responses to the SARS-CoV-2 S-protein and RBD-protein in saliva and sera from SARS-CoV-2 vaccine recipients. The data shown were collated for all vaccine recipients shown in Fig.1A, B and the corresponding panels of ED Fig.1. The longitudinal profiles span a 150-day period before and then ∼60 days after the first immunization (day-0, indicated by the vertical black stippled line). They show saliva and serum IgA, IgG, and IgM antibodies against the S-protein (top two rows, as indicated) and the corresponding reactivities with RBD in saliva (bottom row). Recipients of the Moderna vaccine are represented in red, Pfizer in black.

The longitudinal profiles (Figs. 1A, 1B; ED Fig. 1) and the collated data (Fig. 2) show that the Pfizer and Moderna vaccines rapidly and consistently induce S-protein-specific IgG and IgA in both saliva and sera, with IgM also occasionally detected. Samples were available for 19 individuals at various times during the inter-dose period, with S-protein IgG detectable in saliva from 8 of them during this period (Fig. 1A, B; ED Fig. 1A, B). After their second dose, all 45 vaccine recipients (37 Pfizer, 8 Moderna) were positive for saliva S-protein IgG, with IgA antibodies frequently but not uniformly detected and IgM present only rarely (Table 1). All the vaccine recipients had serum IgG and IgA antibodies after the second dose, and most also had IgM antibodies. Antibody binding to the SARS-CoV-2 RBD in an ELISA was generally quite similar to what was detected using the S-protein (Fig. 2). In contrast, antibodies to the S-protein were not detectable in saliva samples from uninfected control individuals (Fig.1C, ED Fig.1C). Trial participants who became virus-infected before or during the trial did, however, have S-protein IgG antibodies in their saliva, although these infection-elicited responses tended to wane over a multimonth period. IgA antibodies were occasionally detected at lower levels in these saliva samples (Fig. 1D, ED Fig.1D). The observation of S-protein IgG and IgA in the saliva of SARS-CoV-2 infected people is consistent with previous reports (6, 7). When two infected individuals (0001, 0052) were vaccinated, their saliva and serum S-protein IgA and IgG reactivities and, in once case also IgM, were rapidly boosted (Fig. 1D).

The data for saliva and serum S-protein antibodies in Figs. 1 and 2 are derived from ELISAs performed under different conditions of sample dilutions, S-protein coating amounts, etc. (see Methods). In summary, the saliva assay is the more sensitive of the two, and hence the data plots should not be interpreted as indicating there are quantitatively similar IgA and IgG reactivities in the two fluids. To gain an insight into the relative magnitudes of the saliva and serum responses, we titrated a serum sample from an infected individual (D56) and sera from two participants who had each received two doses of the Pfizer vaccine (Cases 0003 and 0007) in the saliva ELISA format, alongside saliva samples from the same vaccine recipients (Fig. 3). Judged by the displacements of the titration curves, we estimate that the titers of S-protein IgA and IgG antibodies present in saliva are ∼1000-fold and ∼10,000 fold lower, respectively, than in serum (Fig. 3). Note that infection serum D56 neutralized SARS-CoV-2 with an ID_50_ titer of 900 in our SARS-CoV-2 pseudovirus-based neutralization assay (8). The much lower S-protein antibody reactivities present in saliva are well below what is required to register in that assay.

**Figure 3.**
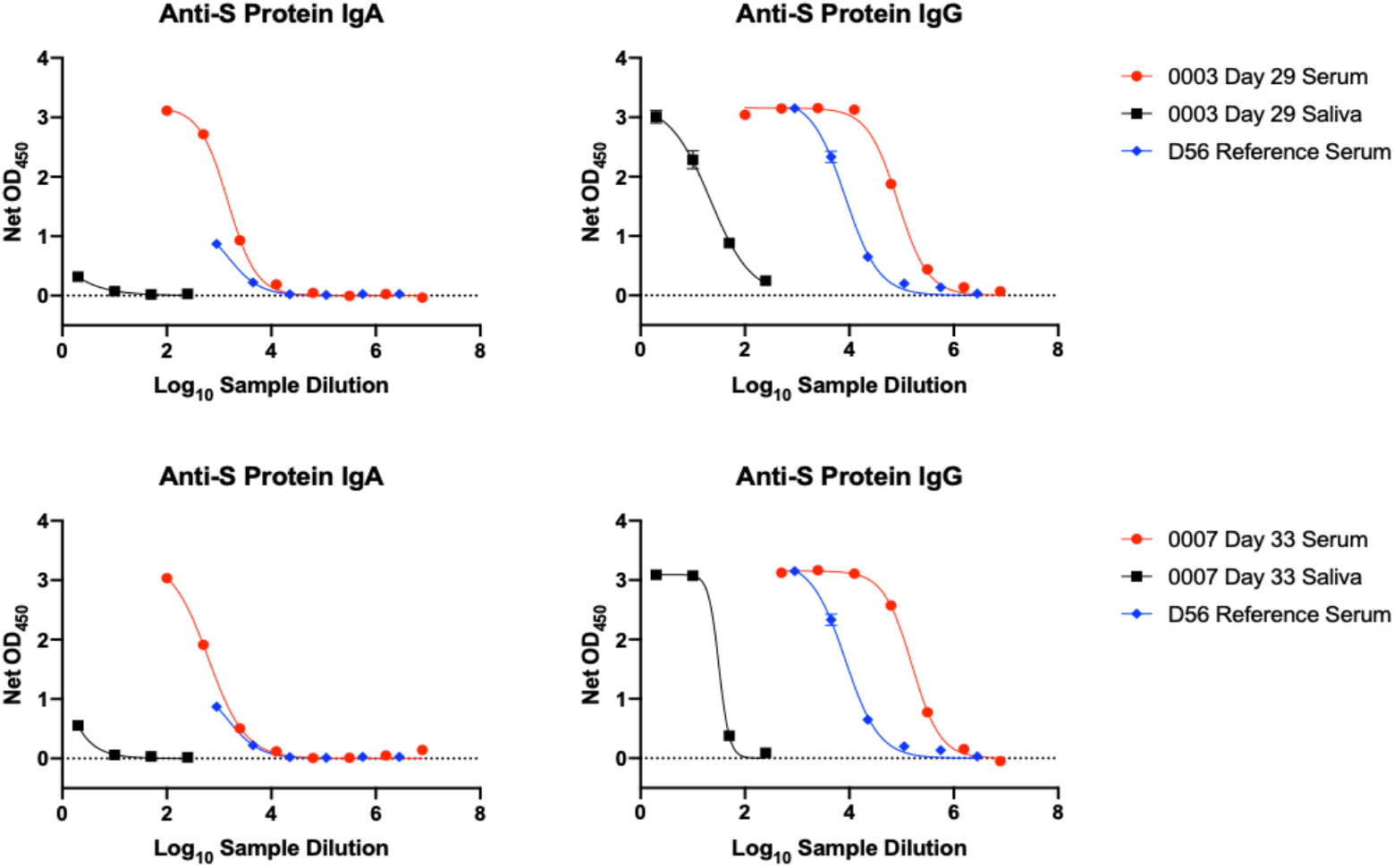
Relative antibody reactivities with S-protein in saliva and sera. A reference serum from a SARS-CoV-2 infected person (D56, not part of the NYP-WELCOME cohort; in blue) and serum (in red) and saliva (in black) samples from two recipients of two doses of the Pfizer vaccine (0003, day-29, top row; 0007, day-33, bottom row) were titrated under the conditions of the ELISA used to detect IgA and IgG in saliva. The displacement of the serum and saliva titration curves suggest that the S-protein IgA and IgG titers in saliva are ∼1000-fold and ∼10,000-fold lower than in sera, respectively.

We show here that the mRNA vaccines induce not just systemic humoral responses that are quantified in serum samples, but also IgG and, to a lesser extent, IgA antibodies to the S-protein and the RBD that are detectable in saliva. The IgA detection antibody we used recognizes both dimeric IgA with and without the secretory component (i.e., SIgA + dIgA). Antibodies in saliva are known to be dominated by SIgA and IgG. Local plasma cells in the stroma of salivary glands secrete dIgA, which traffics through mucosae by interacting with the polymeric Ig receptor. Typically, ∼95% of salivary IgA is in the SIgA form. In contrast, IgG in saliva largely originates from plasma, by transudation from the gingival blood circulation (9-11). IgM is found at lower levels and is also derived from plasma. In healthy people, the total concentrations in saliva are ∼150 µg/ml for SIgA, ∼15 µg/ml for IgG and ∼5 µg/ml for IgM. In contrast, serum concentrations are much higher, ∼1000-fold for IgG and ∼100-fold for total IgA (10, 11). Here, we estimate that the S-protein IgG titers we detected in saliva are ∼10,000-fold lower than in sera, while SIgA titers are ∼1000-fold lower. Accordingly, any SARS-CoV-2 NAbs present in saliva are below assay detection limits. Could vaccine- or infection-induced salivary, and by extension, nasopharyngeal antibodies prevent or limit SARS-CoV-2 infection at the principal portals of entry – the mouth and nose? Our data do not allow this answer to be determined but they do raise the possibility and suggest that additional studies in this area would be fruitful. A recent report of SARS-CoV-2 infection in rhesus macaques shows that virus can replicate in nasal turbinates when it is suppressed in the lungs by passively transferred NAbs (12). Assays of vaccine-induced antibodies extracted from NP swabs might provide additional information. However, the Ig concentrations in tissue sites, e.g., intra-mucosal interstitial fluids, may be impossible to measure by swab-sampling and extraction.

In future studies, it should be possible to determine whether other vaccine designs, such as the Johnson & Johnson adenovirus vector and the Novavax adjuvanted protein, also induce saliva antibodies, and if so to what extent. Comparing the efficacy of these different vaccines at preventing transmission to their abilities to induce mucosal antibody responses may then reveal useful information on any role played by mucosal antibodies.

## Acknowledgements

We thank Dr. Elizabeth Ross, Maria Salpietro (Director of the Institutional Biobank), Ashley Sukhu and Biobank staff members for their contributions to the NYP-WELCOME Cohort. We are grateful to Dr. Prem Lakshamanane (University of North Carolina) and Dr. Andrew Ward (Scripps Research Institute) for the gifts of reagents. We appreciate the administrative support of Kathrina Guemo. RWS is a recipient of a Vici fellowship from the Netherlands Organisation for Scientific Research (NWO). This work was supported by NIH grants P01 AI100652 (JPM) and R01 AI36082 (JPM), and by Weill Cornell Medicine Competitive Research Funding for the NYP-WELCOME trial (SCF).

## Author Contributions

All authors reviewed the manuscript; T.J.K., D.C., S.C.F., P.J.K., and J.P.M. formulated ideas, designed the study and experiments. T.J.K., D.C., E.F., G.D., R.D-T., A.Y., V. C-P, and AC performed experiments and produced reagents; T.J.K., D.C., and P.J.K. analyzed experimental data; E.C, S.C., W.L., Z.Z., P.J.M.B., M.M.C., R.W..S, and S.C.F. provided key reagents. T.J.K., D.C., Z.Z., M.M.C, S.C.F., P.J.K., and J.P.M. supervised work. P.J.K. and J.P.M. wrote the manuscript.

## Competing interests

The authors declare no competing interests.

## Extended Data Figure Legends

**ED Figure 1.**
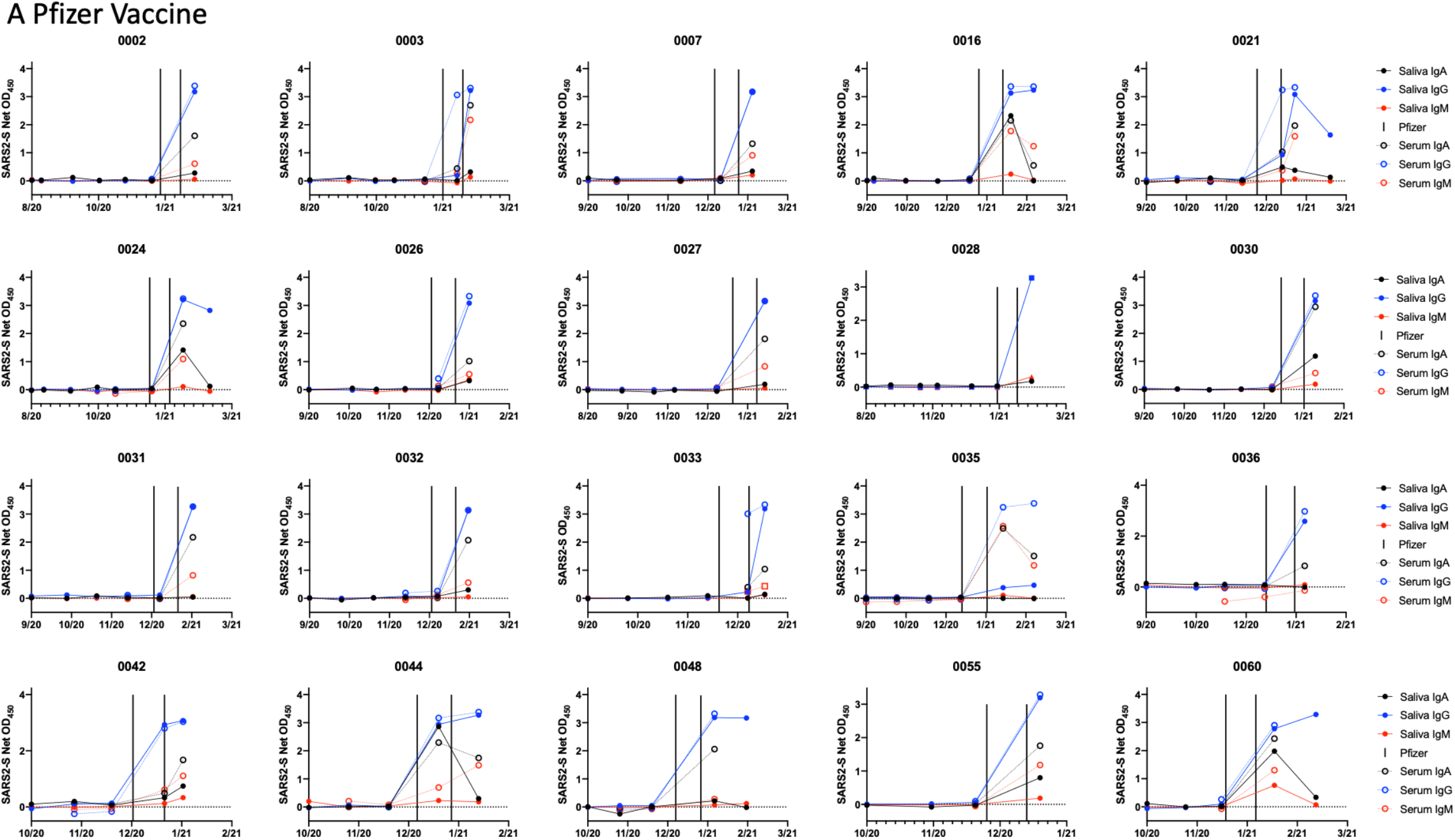
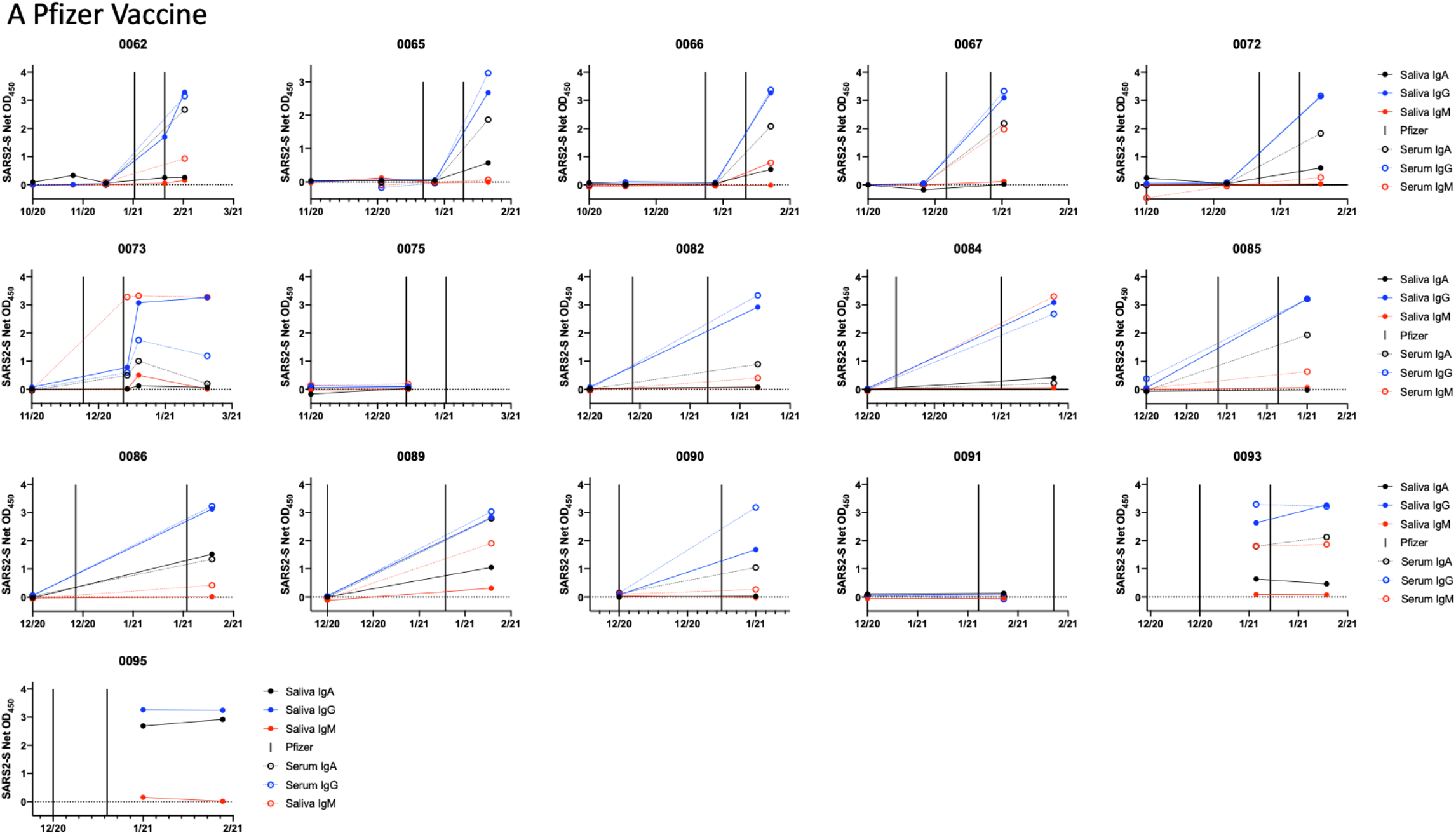
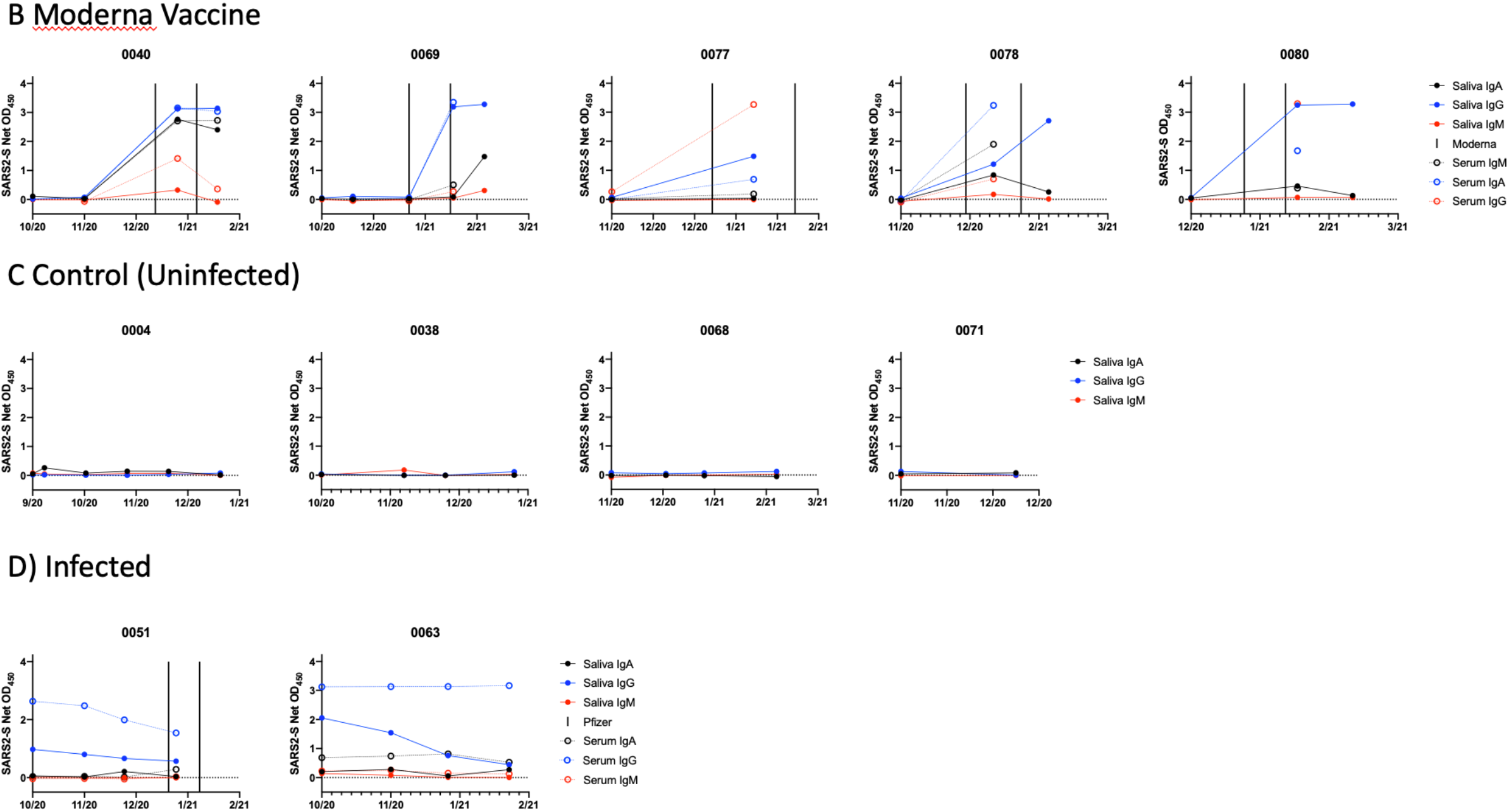
Antibody response to the SARS-CoV-2 S-protein in saliva and sera from SARS-CoV-2 vaccine recipients and infected people. The format of this figure is the same as Figure 1, and shows additional longitudinal profiles for trial participants in Groups 1-4, as indicated.

**ED Figure 2.**
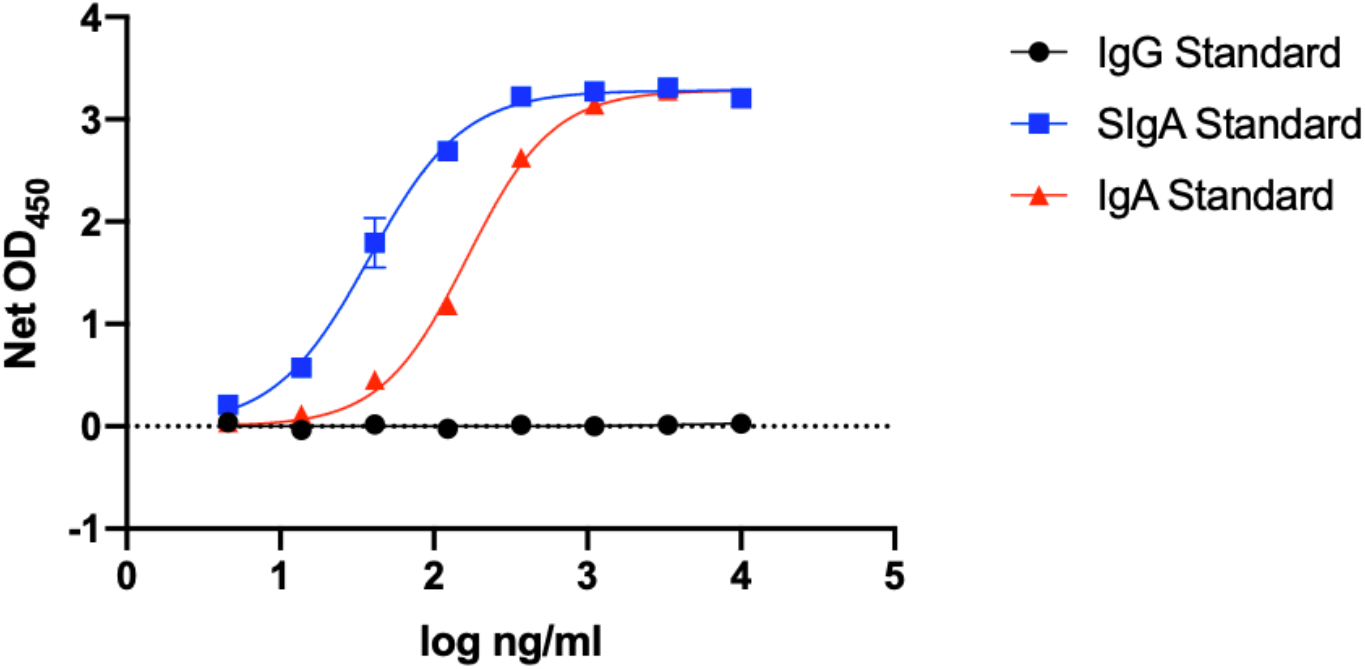
Specificity and sensitivity of IgA detection. Purified human IgG, purified SIgA and recombinant IgA_1_ lambda (see Online Methods) were coated on to ELISA plates and detected with the same goat anti-human IgA-HRP conjugate used in the assays to detect saliva and serum IgA S-protein antibodies. The plot shows net OD_450_ values as a function of the amounts of the three antibodies added to the ELISA wells during the coating stage.

## Online Methods

### The NYP-WELCOME cohort

The NYP-WELCOME (WEilL COrnell Medicine Employees) trial was initiated on June 16th, 2020 at Weill Cornell Medicine (WCM) and New York Presbyterian Hospital (NYP). The aim of the trial is to study the determinants of the diverse COVID-19 outcomes (from infection to resolution or death and before, during and after vaccination) among an exposed population in Manhattan, consisting of NYP-Weill Cornell Medicine asymptomatic healthcare workers who volunteered to participate. The study is designed to prospectively collect sequential specimens and questionnaires twice a month for first 3 months and then monthly for 2 years, during the Covid-19 pandemic. To date (3/9/21), 97 participants have completed 527 study visits. Two participants became SARS-CoV-2-infected during the trial, as determined by a positive nasopharyngeal RT-PCR test. Six other participants were infected prior to joining the trial, as judged by their positivity for serum S-protein antibodies at multiple time-points. Of the six infected people, three were vaccinated. The Pfizer and Moderna mRNA vaccines became available at NYP-WCM after December 15th, 2020, and 78/97 participants have now received at least one dose of either vaccine (Pfizer, n = 63 participants; Moderna, n = 15). Samples derived from the NYP-WELCOME trial are stored in an institutional biorepository that now contains ∼11,300 specimens including PBMCs, saliva, stool, urine, plasma, serum and nasopharyngeal (NP) swab extracts. In this study, we used serum and saliva samples only. The WCM Institutional Review Board approved the study on 6.3.2020 with the protocol number 20-04021831 and title WELCOME trial (Weill Cornell Medicine Employees) with Dr. Silvia C. Formenti MD as the Principal Investigator.

### Sample collection and processing

Saliva and blood were collected from members of the NYP-WELCOME cohort on an initially biweekly and then monthly basis. As the trial was not established as a vaccine study, there was no coordination between the dates of vaccine administration and sample collection. Blood was processed into serum that was heat-inactivated at 56°C. Nonidet-P40 (NP-40) non-ionic detergent was added to saliva samples to a final concentration of 0.05% (vol/vol), both to inactivate any SARS-CoV-2 present and as a preservative. Saliva extracts were passed through a 22 µm filter under sterile conditions, before addition of protease inhibitors to inhibit sample degradation. The sources and final concentrations of these inhibitors were as follows: Aprotinin, 8.5 µg/ml (Sigma Aldrich); phenylmethanesulfonyl fluoride, 5.9 mM (Sigma-Aldrich 93482); sodium orthovanadate, 1.2 mM (Sigma-Aldrich S6508). The processed saliva samples were stored at −20°C.

### S-proteins and RBD-proteins

The expression construct for the pre-fusion S2-P stabilized SARS-CoV-2 S-Foldon-StreptagII S-protein ectodomain used in this study was derived as follows. The gene encoding residues 1-1138 from the Wuhan-Hu-1 strain (Genbank MN908947.3) was modified by introducing proline substitutions at amino acid positions 986 and 987 and a “GGGG” substitution at amino acids 682-685 (the furin cleavage site). The modified gene was then cloned into a pPPI4 plasmid containing a T4 trimerization domain followed by a Strep-tag® II (13). The SARS-CoV-2-RBD-StrepII expression construct was a gift from Dr. Prem Lakshamanane (University of North Carolina) and has been described previously (14). The constructs were expressed transiently in ExpiCHO cells. Briefly, cells were transfected at a density of 6 x 10^6^ per ml by adding FectoPRO (Polyplus-transfection SA) transfection reagents. For a 1 L transfection 800 µg of plasmid, 1.6 ml of transfection reagent and 500 µl of FectoPRO booster were added to 40 ml of Opti-MEM (Thermo Fisher Scientific). Culture supernatants were harvested 3 days post-transfection, centrifuged for 1 h at 6000 rpm and passed through a 0.2 μm filter (Thermo Fisher Scientific). BioLock-Biotin blocking solution (IBA Lifesciences) was then added before the treated supernatants were passed over StrepTactin™ Sepharose resin (GE Healthcare). The S-protein or RBD was eluted in 2.5 mM desthiobiotin in 100 mM Tris-Hcl, 150 mM NaCl, 1mM EDTA, pH 8.0. Proteins were dialyzed into PBS and concentrated using Vivaspin protein spin columns with a 100 kDa (S-protein) or 10 kDa (RBD protein) molecular weight cutoff (GE Healthcare). Protein concentrations were determined using the BCA protein assay kit (Thermo Fisher) and purity was assessed using a 4 to 16%, Bis-Tris Native PAGE gel system (Invitrogen). For SDS-PAGE, purified proteins were denatured with 0.1 mM dithiothreitol (DTT) before loading onto a 4 to12% Bis-Tris Gel NuPAGE gel (Invitrogen).

### ELISA procedures

The assay used to quantify S-protein and anti-RBD antibodies was modified from what was described previously (15). For work with serum samples, S-proteins or RBD proteins (200 ng in 100 µl) were coated overnight onto 96-well plates at 4°C. After 3 washes with PBS/0.05% Tween-20 (PBST), the wells were blocked for 1 h with 4% (w/vol) powdered milk/PBS (150 µl/well). Serum was initially diluted 1/100 in PBS containing 4% milk and 20% sheep serum, and serially diluted as needed, and added to the wells for 1 h. Bound antibodies were detected using goat anti-human horse radish peroxidase (HRP)-conjugated antibodies: anti-IgA from Southern Biotech (2050-05), diluted 1/3000 in 4% milk/PBS; anti-IgG from Jackson ImmunoResearch (109-035-008), diluted 1/5000; anti-IgM from Jackson ImmunoResearch (109-035-043), diluted 1/5000. After washing, a 50 µl volume of HRP substrate (Thermo Scientific 34029) was added to each well for 3 min before color development was terminated with 0.3 N sulfuric acid. The plates were read at 450 nm using an Enspire instrument (Perkin Elmer).

For work with saliva extracts, the following modifications were made to increase assay sensitivity and conserve sample volume. The 96-well plates were replaced by 384-well plates (Thermo Scientific 464718) and the incubation volumes were correspondingly smaller (10 µl/well) except the blocking buffer (100 µl/well). The amount of S-protein or RBD protein added to the wells was 100 ng in 10 µl, to create a higher coating density. The HRP substrate volume was increased to 25 µl/well and the reaction time lengthened to 15 min. Accordingly, the colorimetric signals derived from the saliva and NP assays are not directly comparable to those from the serum assay (see Results).

S-IgA was detected as follows: half-area plates (Corning 3690) were coated overnight with goat anti-human IgA (Bio Rad STAR141) (250 ng in 50 µl), washed 3 times with PBST and blocked with 75 µl of 2% (w/vol) powdered milk/PBS for 1 h, and then washed 3 more times with PBST. Saliva was initially diluted 1/4 in PBS containing 20% sheep serum and 2% powdered milk and then serially diluted as needed. A SIgA standard (Bio Rad Purified Human Secretory IgA PHP133) was similarly serially diluted from an initial stock at 5 µg/ml. The saliva or SIgA samples were added to wells for 1 h at room temperature, the plates were washed 3 times with PBST before addition of 50 µl of a 1/500 dilution of Monoclonal Anti-Secretory Component IgA (Sigma I6635) that had been conjugated to HRP using the Conjugation Kit-Lighting-Link according to the manufacturer’s instructions (ab102890 HRP). After incubation for 1 h at room temperature, the plates were again washed 3 times with PBST before addition of HRP substrate (50 µl). The colorimetric reactions were terminated and the plates read as described above.

To assess the specificity and sensitivity of the IgA detection procedure, purified human IgG (Sigma 14506), purified Secretory IgA (SIgA) (BioRad PHP133) or recombinant IgA_1_ lambda (BioRad HCA172) were coated onto ELISA plates (Corning 3690) and probed with goat anti-Human IgA-HRP (Southern Biotech 2050-05). The assay detected both IgA and SIgA (ED Fig. 2). The IgA-detection conjugate is specific for IgA and does not cross-react with IgG. Both SIgA and recombinant IgA standards are detected. The ∼5-fold more sensitive detection of SIgA could arise from coating efficiency differences *vs*. IgA under the ELISA conditions used and/or the properties of the conjugate (ED Fig. 2).

### Data analysis

ELISA-derived net OD_450_ values for 4-fold dilutions of saliva and 100-fold dilutions of sera were determined by subtracting background values obtained from wells containing no S-protein or RBD. The cut-off for positive antibody detection was set to 0.300 net OD_450_, corresponding to 6-times the average OD_450_ value derived from the negative samples. The net OD_450_ values were plotted longitudinally for each study participant.

## Notes

### Competing Interest Statement

The authors have declared no competing interest.

